# Exploring the microbial community structure and metabolic gene clusters during silage fermentation of paper mulberry, and developing the high-protein woody plant as ruminant feed

**DOI:** 10.1101/2020.07.08.194605

**Authors:** Zhumei Du, Lin Sun, Chao Chen, Jing Lin, Fuyu Yang, Yimin Cai

## Abstract

To develop a new high-protein woody forage resource for livestock, we applied PacBio single-molecule, real-time (SMRT) sequencing technology to explore the community structure, species diversity, and metabolic gene clusters of nature microbes associated with paper mulberry (PM) silage fermentation. The microbial diversity and abundance were rich in PM raw material and decreases with the progress of silage fermentation. Woody ensiling is process that the dominant bacteria shifted from Gram-negative pathogenic *Proteobacteria* to Gram-positive beneficial *Firmicutes*. Lactic acid bacteria became the most dominant bacteria that affected fermentation quality in the terminal silages. Global and overview maps, carbohydrate metabolism and amino acid metabolism were the important microbial metabolic pathways that impact final fermentation product of silage. PM is rich in nutrients and preserved well during ensiling, indicating PM can develop as new woody resources suitable for ruminants. PacBio SMRT sequencing revealed specific microbial-related information about silage.

**IMPORTANCE:** In the tropics, there is often a shortage of forage during the dry season. Failure to obtain high-quality feed will reduce the milk and meat production of ruminants. Therefore, it is essential to maximize the use of land and biomass resources through strategic development of alternative feed. Paper mulberry (PM) is a perennial deciduous tree in tropics, with a variety of nutrients and biologically active ingredients, and it adapts to various soils and climates, with high production capacity, and low cultivation costs. In order to develop new potential woody forage, we firstly used PacBio single-molecule real-time (SMRT) sequencing technology to explore the community structure, species diversity and metabolic gene clusters of natural microorganisms related to the fermentation of silage. PacBio SMRT revealed information about specific microorganisms related to silage, indicating PM can prepare as good-quality silage, and will become a new potential woody feed resources for livestock.

---

In tropical regions, livestock farming often exceeds forage availability, especially in the dry season, leading to pasture degradation (1). The most important factor limiting livestock productivity is the seasonality of food supply in terms of pasture quantity and quality (2). Generally, less forage is produced in the dry season. Limited access to high-quality roughage reduces milk and meat production in ruminants (2), which can damage the health and welfare of animals and increase the incidence of diseases (3). Therefore, it is critical to maximize the use of land and biomass resources to ensure satisfactory animal growth index by controlling livestock populations, supplementation with alternative feed, and strategic development of forage resources.

In recent years, nutrient-enriched woody plants as new forage resources have been applied to cope with the challenges posed by feed shortages and rapid development of the livestock industry (4). Paper mulberry [PM, *Broussonetia papyrifera* (L.) Vent.] is considered a representative of such alternative, woody plants to supplement ruminant feed. PM is a perennial deciduous tree or shrub from the Moraceae family that is native to eastern Asia, which has high nutritional value, a variety of biologically active ingredients, high production capacity, and low cultivation and production costs. Moreover, it is adapted to various soils and climates; therefore, it is economically significant for local animal production.

In the tropics, PM is harvest mainly during the summer rainy season, which characterized by high temperatures, heavy rainfall, and humidity levels as high as 80%, such that traditional hay production methods may not be suitable for PM in this region. Preparation and preservation of PM silage is among the most effective techniques to overcome the gap between annual livestock production and the seasonal imbalance in available forage.

Microbial additives, including lactic acid bacteria (LAB) inoculant and cellulase enzymes play an important role in improving silage fermentation and nutrient utilization for ruminants (2). Adjustment of the moisture content, water-soluble carbohydrate (WSC) content, and epiphytic LAB of forage also directly influences bacterial activity during the fermentation stage. Recently, next-generation sequencing has been applied for quantitative analysis of the silage microbiome (5). Third-generation Pacific Biosciences (PacBio) single-molecule real-time (SMRT) sequencing technology generates long sequence reads that improve the sensitivity and accuracy of classification and allow for relatively high taxonomic resolution to the species level (6, 7). Several studies have evaluated the efficacy of feed additives and wilting to improve the fermentation quality of forage crops and grasses silage.

However, little information on woody plant silage is available. To develop new high-protein woody forage resources for animal production, we used PacBio SMRT technology to explore the community structure, species diversity, and metabolic gene clusters of natural microbes associated with PM silage fermentation.

## RESULTS

### The bacterial community and diversity of PM and WPM before and after ensiling

Based on SMRT sequencing of the full-length 16S rRNA gene of silage bacteria, the number of reads for 10 samples ranged from 5,426 to 10,385, with an average of 8,898. Prior to ensiling, the ACE, Chao1, Shannon, and Simpson indexes were lower (*P* < 0.05) in PM than those in wilt PM (WPM) (Table 1). After ensiling, the CH (LAB inoculant Chikusou-1, *Lactobacillus plantarum*, Snow Brand Seed Co., Ltd, Sapporo, Japan) inoculated PM silages produced lower Shannon and Simpson indexes than control silages, but the LAB inoculants increased the microbial diversity of WPM than that in control. A total of 2,926 operational taxonomic units (OTU) were clustered at the 3% dissimilarity level. The OTUs showed trends similar to the trend of the Simpson and Shannon indexes. Higher numbers of bacterial core microbiome OTUs were identified in PM and WPM (Fig. 1A) than after ensiling (Fig. 1B). The bacterial core microbiomes of final PM and WPM raw material were composed of 424 shared OTUs and 161 and 150 unique OTUs for WPM and PM, respectively. The number of unique OTUs decreased sharply to 116 for the PM control silage and 126 for the WPM control silage. The unique OTUs ranged from 14 to 66 for PM silages (Fig. 1C) and from 17 to 50 for WPM silages (Fig. 1D).

**TABLE 1.** General information of sequence and bacterial diversity

| Sample ID | Number of reads | ACE | Chao1 index | Shannon index | Simpson index | Coverage |
| --- | --- | --- | --- | --- | --- | --- |
| PM | 10138 | 192.12 | 210.25 | 0.10 | 3.37 | 0.9936 |
| WPM | 8133 | 199.64 | 218.26 | 0.13 | 3.44 | 0.9927 |
| PM silage |  |  |  |  |  |  |
| Control | 8611 | 289.53 | 318.64 | 0.08 | 3.04 | 0.9471 |
| CH | 9874 | 216.60 | 267.26 | 0.06 | 2.76 | 0.9946 |
| AC | 9441 | 284.71 | 378.75 | 0.08 | 3.02 | 0.9349 |
| CH+AC | 8789 | 284.85 | 377.38 | 0.09 | 3.01 | 0.9776 |
| WPM silage |  |  |  |  |  |  |
| Control | 5426 | 285.86 | 293.78 | 0.10 | 3.11 | 0.9772 |
| CH | 10385 | 305.70 | 487.90 | 0.12 | 3.38 | 0.983 |
| AC | 9134 | 299.59 | 485.75 | 0.12 | 3.40 | 0.9867 |
| CH+AC | 9047 | 290.25 | 438.23 | 0.11 | 3.37 | 0.9808 |

**FIG 1.**
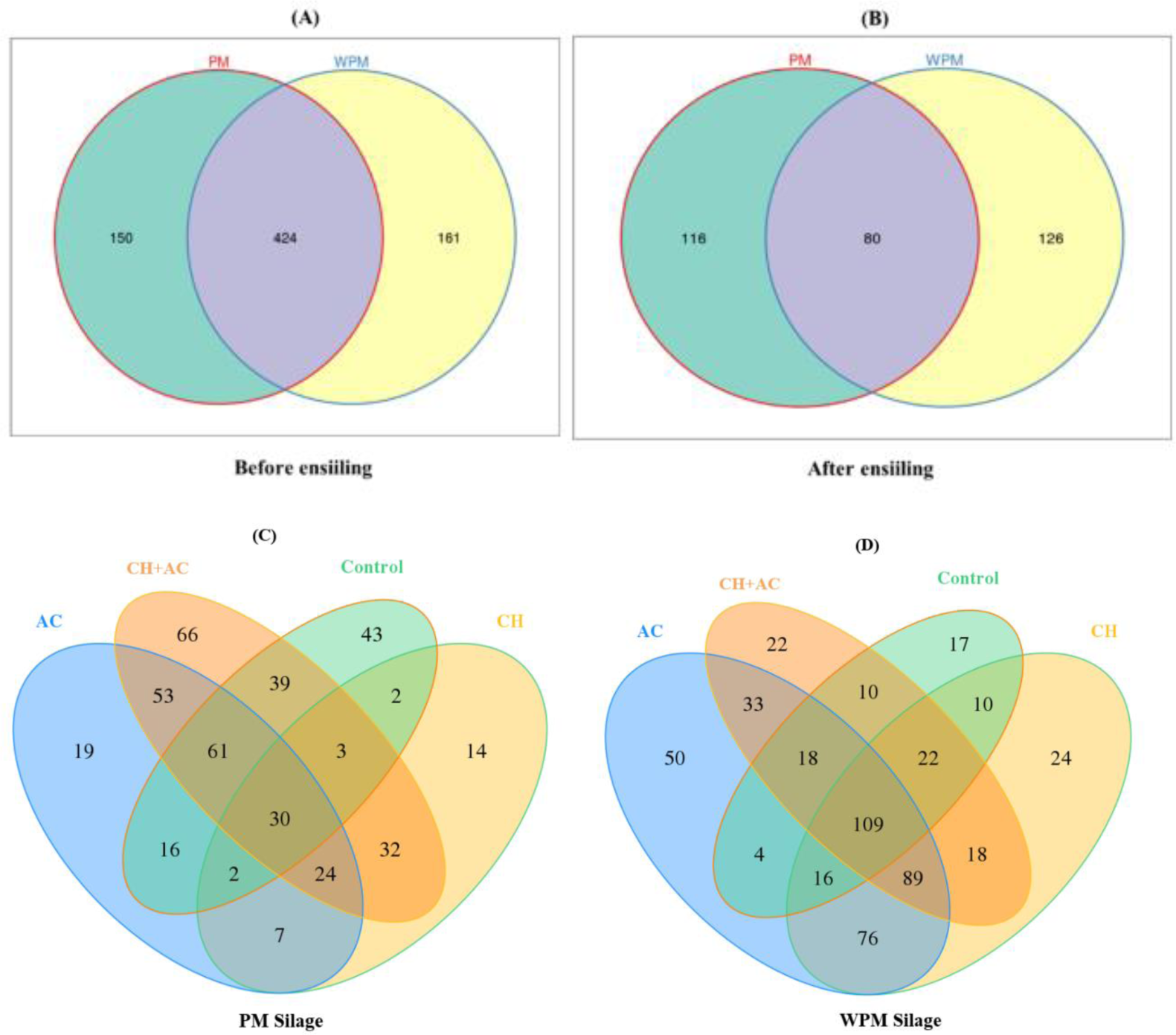
Venn diagram depicting unique or shared bacterial OTUs (97% sequence identity) in PM and WPM before and after ensiling. The number of bacterial core microbiome OTUs is highlighted with a bold black frame: (A) The number of OTUs shared by PM and WPM samples before ensiling; (B) The number of OTUs shared by PM and WPM control silages. (C) The number of OTUs shared by all the PM silages; (D) The number of OTUs shared by all the WPM silages. OTU, operational taxonomic unit; PM, paper mulberry; WPM, wilted paper mulberry; CH, *Lactobacillus plantarum*; AC, *Acremonium cellulase*; CH + AC, combination of CH and AC.

### The relative abundance of bacteria at genus level in PM and WPM before and after ensiling

From the Fig. 2A and B, it emphasized that the main dominant bacterial genus were *Pseudomonas* (42.32%, percentage of total bacteria), *Pantoea* (26.26%), *Serratia* (10.06%), *Erwinia* (6.67%), and *Rahnella* (0.85%) in the PM raw material. The main dominant genus in the WPM raw material were *Acinetobacter* (49.43%), *Pantoea* (16.30%), *Aerococcus* (6.87%), *Enterobacter* (6.17%), and *Staphylococcus* (4.53%). After fermentation, *Lactobacillus* was found to be the predominant genus for all the silages expect for CH + AC (cellulase enzyme, *Acremonium cellulase*, Meiji Seika Pharma Co., Ltd, Tokyo, Japan) of WPM. The relative abundance of *Lactobacillus* ranged from 72–98% and 40–61% in PM and WPM silages, respectively. They were followed by *Pediococcus* for PM, and *Enterobacter* for WPM. The highest abundance of *Lactobacillus* was observed in in the CH-treated PM silage.

**FIG 2.**
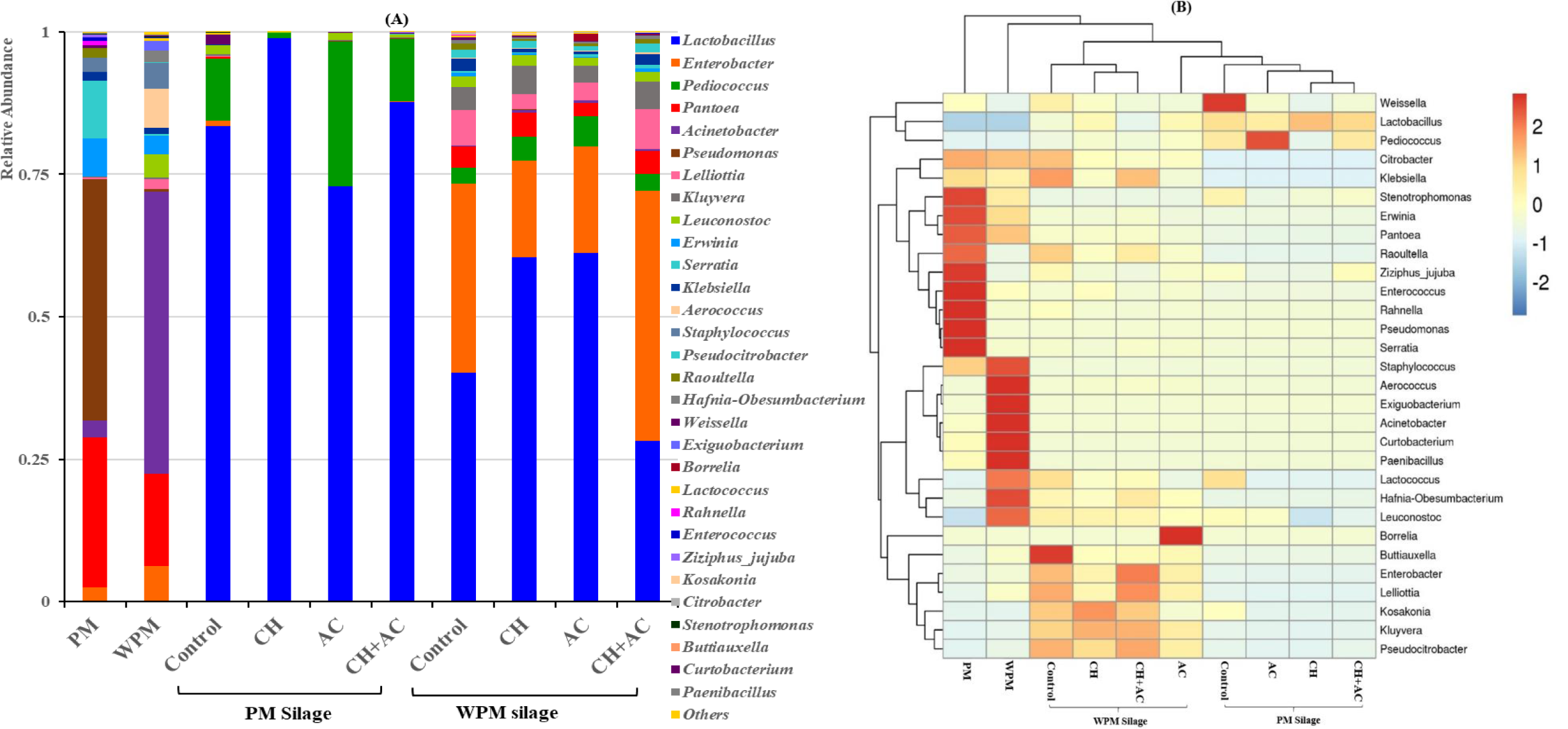
Bacteria community (A), hierarchical clustering and heat map analyses (B) in the PM and WPM silages at genus levels before and after ensiling. PM, paper mulberry; WPM, wilted paper mulberry. CH, *Lactobacillus plantarum*; AC, *Acremonium cellulase*; CH + AC, combination of CH and AC. For the Fig. 2B, X-axis represents sample name, Y-axis represents the genus. Block with red color indicates the high abundance of the genus, the blue block indicates the low abundance of the genus. The absolute value of ‘Z’ represents the distance between the raw score and the mean population of the standard deviation. The ‘Z’ is negative when the raw score is below the mean, and vice versa. The hierarchical clustering tree appears on the upside of block while the phylogenetic tree appears on the left side of block.

### The relative abundance of bacteria at species level in PM and WPM before and after ensiling

The bacterial communities of silages bacteria at the species level (analyzed by SMRT) before and after 60 days of ensiling are shown in Fig. 3A and B. *Pantoea agglomerans* (22.99%, percentage of total bacteria), *Pseudomonas sp.* (19.32%), *Pseudomonas koreensis* (14.07%), and *Serratia liquefaciens* (7.03%), and *Pseudomonas coleopterorum* (7.01%) were the dominant species in the PM, but the main species in the WPM were *Acinetobacter sp.* (48.45%), *Pantoea agglomerans* (12.02%), *Enterobacter sp.* (7.04%), and *Streptomyces alboniger* (6.72%) of the total bacteria, respectively, whereas the proportion of *Lactobacillus plantarum* was almost undetectable in the PM (0.02%) and WPM (0.01%) raw material. After fermentation, the most abundant microbe species in the control PM silage was *Lactobacillales bacterium* (16.76%), followed by *Lactobacillus vaccinostercus* (16.40%) and *Lactobacillus plantarum* (15.93%). For the control treatment of WPM silages, the most abundant species was *Enterobacter sp*. (34.69%), followed by *Lactobacillus sp.* (20.19%), *Lelliottia amnigena* (2.08%), and *Klebsiella oxytoca* (1.25%).

**FIG 3.**
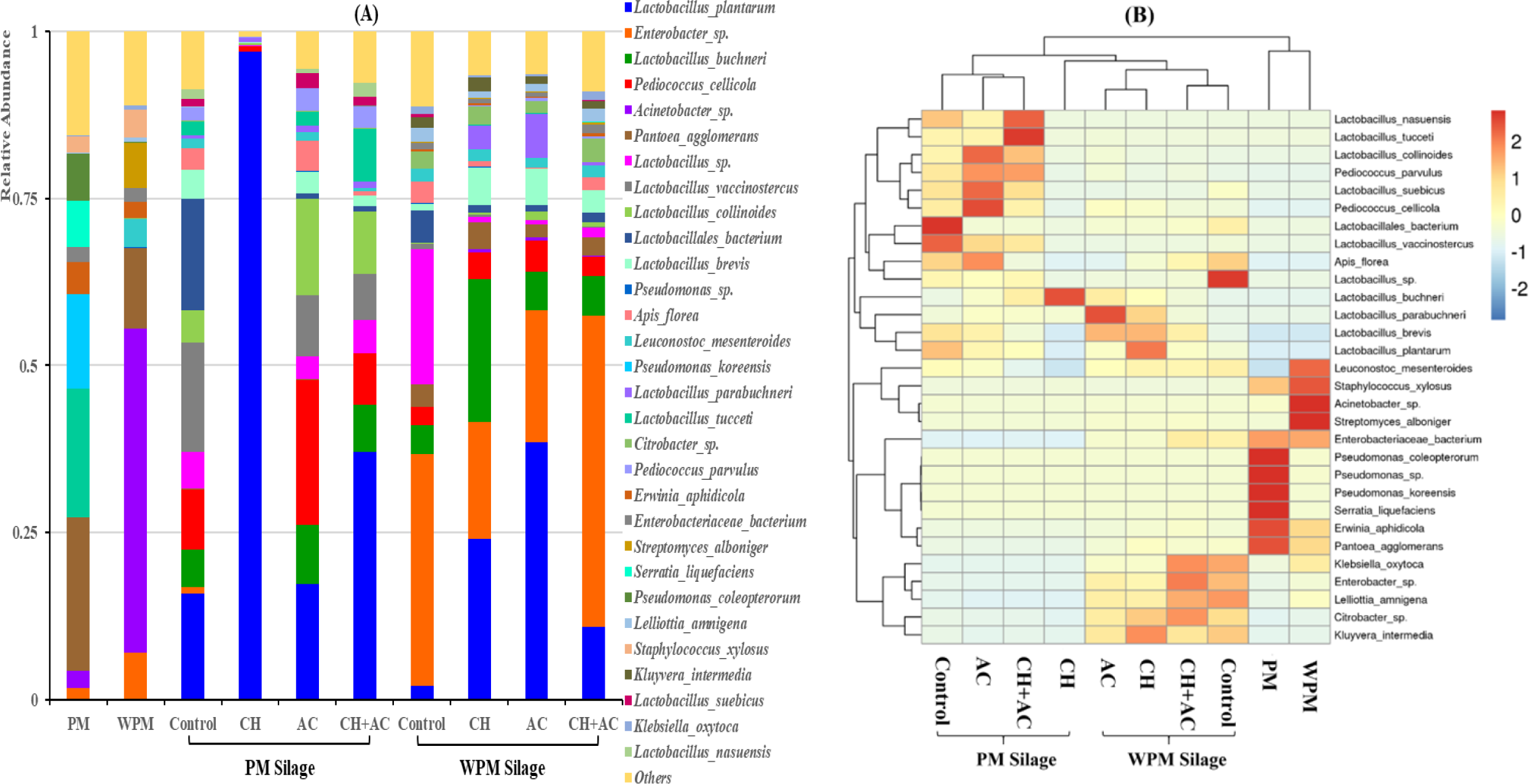
Bacteria community (A), hierarchical clustering and heat map analyses (B) in the PM and WPM silages at species levels before and after ensiling. PM, paper mulberry; WPM, wilted paper mulberry; CH, *Lactobacillus plantarum*; AC, *Acremonium cellulase*; CH + AC, combination of CH and AC.

### Phylogenetic circle tree at species level of PM and WPM before and after ensiling

Based on phylogenetic circle tree of 16S rRNA, there were three types phylums in PM (Fig. 4A) and WPM (Fig. 4B) raw material, there were three types phylums, they were *Firmicutes, Actinobacteria*, and *Proteobacteria*, respectively. *Proteobacteria* was the most abundant phylum in the samples, on average accounting for 96.33 and 81.47% in PM and WPM, respectively, followed by *Firmicutes* with an average of 3.48 and 18.08%, *Actinobacteria* with an average of 0.19 and 0.44%. After ensiling, *Firmicutes* was the most abundant phylum in the samples, on average accounting for 97.92 and 45.91% in PM (Fig. 4C) and WPM (Fig. 4D), respectively, followed by *Proteobacteria* with an average of 2.08 and 54.09%.

**FIG 4.**
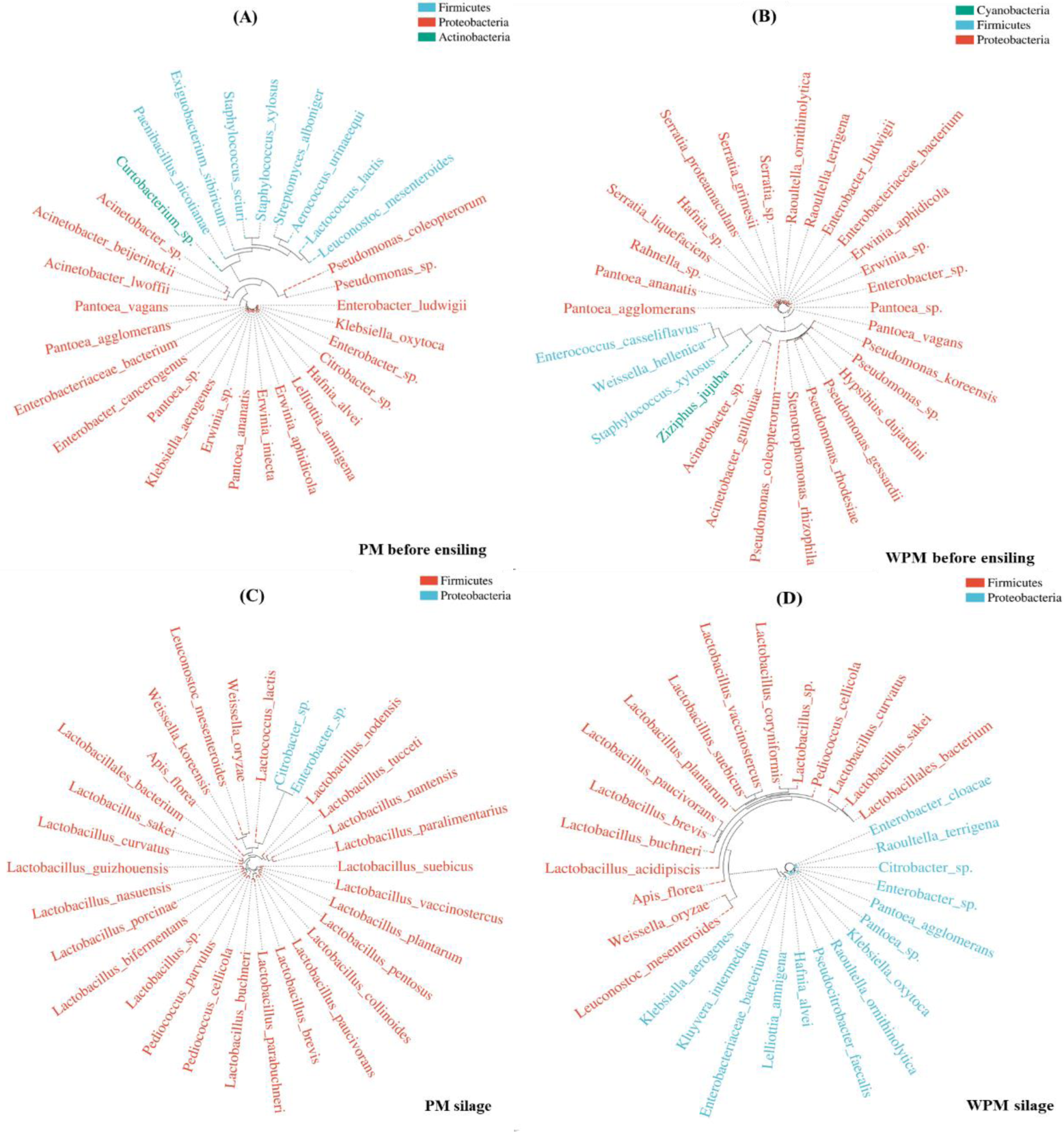
Phylogenetic circle tree based on sequences representative of the OTUs at species level of PM and WPM before and after ensiling. OTU, operational taxonomic unit; PM, paper mulberry; WPM, wilted paper mulberry.

### Chemical composition and microbial population of PM and WPM before ensiling

The pH, lactic acid buffer capacity (LBC), microbial population, chemical composition, protein, energy, in vitro dry matter digestibility (IVDMD), gas production, and macro mineral of PM and WPM before ensiling are shown in Table 2. The LBC of PM was 1241.07 mEq/kg of dry matter (DM), which value of WPM was lower than the PM samples (*P* < 0.05), while the pH was opposite. Prior to ensiling, the DM of PM was 20.05%, and increased by 26.08% in WPM. The organic matter (OM), crude protein (CP), ether extract (EE), neutral detergent fiber (NDF), acid detergent fiber (ADF), acid detergent lignin (ADL), and WSC were 89.79, 29.94, 4.10, 39.17, 19.84, 6.23, and 8.40% of DM, respectively, for PM. The chemical composition of PM exhibited no major changes during the wilting process. The low counts of LAB (10^4^ cfu/g of fresh matter (FM)) and higher amounts of undesirable microorganisms (10^3^–10^5^ cfu/g of FM) were observed in the PM and WPM raw material.

**TABLE 2.**
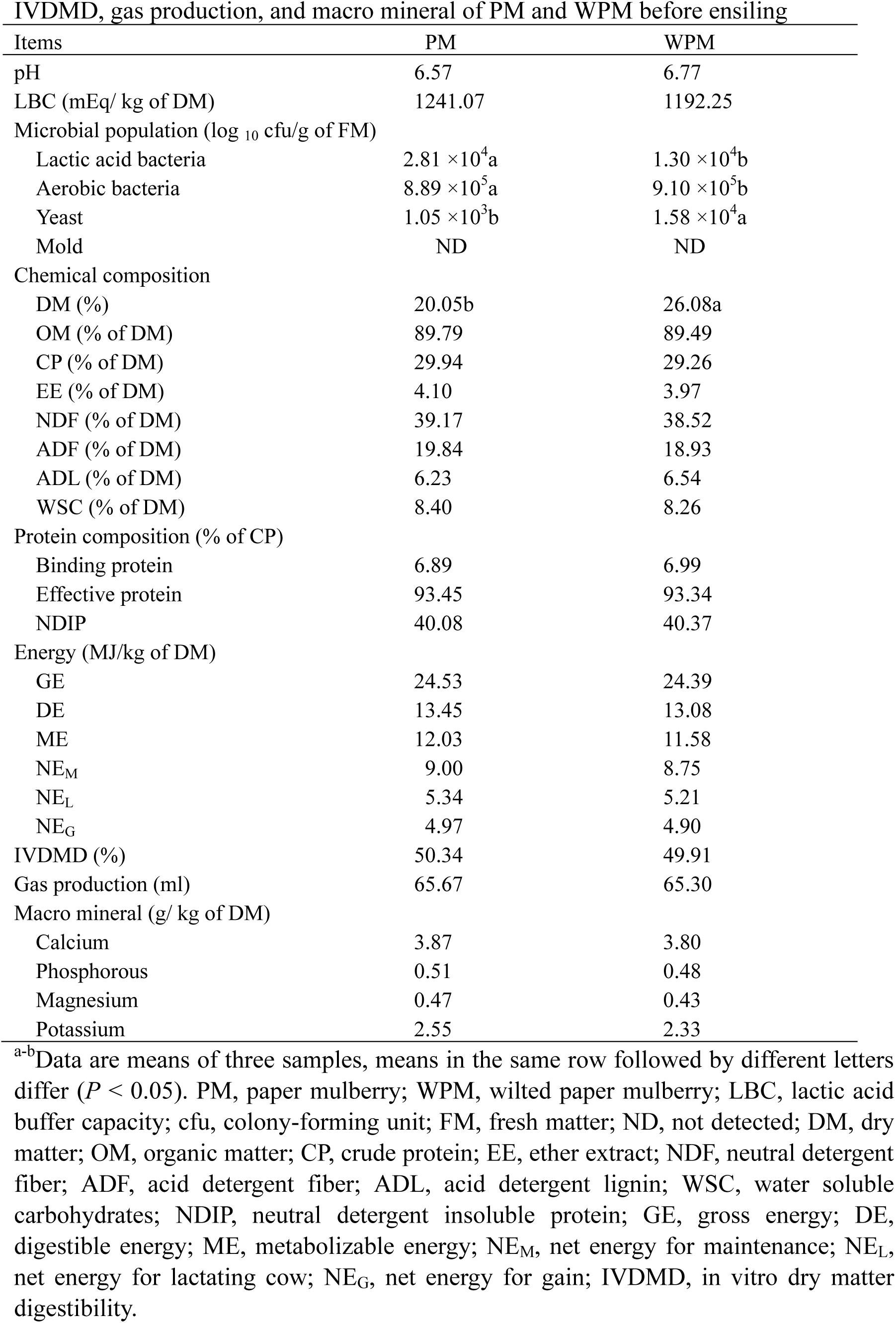
pH, LBC, microbial population, chemical composition, protein, energy, IVDMD, gas production, and macro mineral of PM and WPM before ensiling

| Items | PM | WPM |
| --- | --- | --- |
| pH | 6.57 | 6.77 |
| LBC (mEq/ kg of DM) | 1241.07 | 1192.25 |
| Microbial population (log<sub>10</sub> cfu/g of FM) |  |  |
| Lactic acid bacteria | 2.81 × 10<sup>4</sup>a | 1.30 × 10<sup>4</sup>b |
| Aerobic bacteria | 8.89 × 10<sup>5</sup>a | 9.10 × 10<sup>5</sup>b |
| Yeast | 1.05 × 10<sup>3</sup>b | 1.58 × 10<sup>4</sup>a |
| Mold | ND | ND |
| Chemical composition |  |  |
| DM (%) | 20.05b | 26.08a |
| OM (% of DM) | 89.79 | 89.49 |
| CP (% of DM) | 29.94 | 29.26 |
| EE (% of DM) | 4.10 | 3.97 |
| NDF (% of DM) | 39.17 | 38.52 |
| ADF (% of DM) | 19.84 | 18.93 |
| ADL (% of DM) | 6.23 | 6.54 |
| WSC (% of DM) | 8.40 | 8.26 |
| Protein composition (% of CP) |  |  |
| Binding protein | 6.89 | 6.99 |
| Effective protein | 93.45 | 93.34 |
| NDIP | 40.08 | 40.37 |
| Energy (MJ/kg of DM) |  |  |
| GE | 24.53 | 24.39 |
| DE | 13.45 | 13.08 |
| ME | 12.03 | 11.58 |
| NE<sub>M</sub> | 9.00 | 8.75 |
| NE<sub>L</sub> | 5.34 | 5.21 |
| NE<sub>G</sub> | 4.97 | 4.90 |
| IVDMD (%) | 50.34 | 49.91 |
| Gas production (ml) | 65.67 | 65.30 |
| Macro mineral (g/ kg of DM) |  |  |
| Calcium | 3.87 | 3.80 |
| Phosphorous | 0.51 | 0.48 |
| Magnesium | 0.47 | 0.43 |
| Potassium | 2.55 | 2.33 |

<sup>a-b</sup>Data are means of three samples, means in the same row followed by different letters differ ( $P < 0.05$ ). PM, paper mulberry; WPM, wilted paper mulberry; LBC, lactic acid buffer capacity; cfu, colony-forming unit; FM, fresh matter; ND, not detected; DM, dry matter; OM, organic matter; CP, crude protein; EE, ether extract; NDF, neutral detergent fiber; ADF, acid detergent fiber; ADL, acid detergent lignin; WSC, water soluble carbohydrates; NDIP, neutral detergent insoluble protein; GE, gross energy; DE, digestible energy; ME, metabolizable energy; NE<sub>M</sub>, net energy for maintenance; NE<sub>L</sub>, net energy for lactating cow; NE<sub>G</sub>, net energy for gain; IVDMD, in vitro dry matter digestibility.

### Microbial population, fermentation quality, and chemical composition of silages

The microbial population, fermentation quality, chemical composition, IVDMD, and gas production of silages are shown in Table 3. The LAB counts were 10^6^ to 10^7^ cfu/g of FM in all the PM silages and 10^4^ in all the WPM silages. The aerobic bacteria and yeast counts in all silages were 10^4^ except for CH + AC (a mixture of LAB and cellulase) in the WPM silage (10^3^). Meanwhile, coliform bacteria and molds were below detectable levels (< 10^2^) in all silages. In the CH, AC, and CH + AC-treated silages, the LAB count was significantly (*P* < 0.05) higher, while their aerobic bacteria was significantly (*P* < 0.05) lower than that of the control. The silage (S), additive (A), and their interaction between S and A (S × A) significantly (*P* < 0.05) influenced the LAB, aerobic bacteria, and yeast counts.

**TABLE 3.** Microbial population, fermentation quality, chemical composition, IVDMD, and gas production of PM and WPM silages after 60 days of ensiling

| Item | PM silage |  |  |  | WPM silage |  |  |  | *P*-value |  |  |
| --- | --- | --- | --- | --- | --- | --- | --- | --- | --- | --- | --- |
|  | Control | CH | AC | CH + AC | Control | CH | AC | CH + AC | S | A | S × A |
| Microbial population (log<sub>10</sub> cfu/g of FM) |  |  |  |  |  |  |  |  |  |  |  |
| Lactic acid bacteria | 6.85d | 7.98a | 7.00c | 7.52b | 4.20c | 4.37b | 4.58a | 4.16d | * | ** | * |
| Coliform bacteria | ND | ND | ND | ND | ND | ND | ND | ND | ND | ND | ND |
| Aerobic bacteria | 4.55a | 4.20c | 4.41b | 4.04d | 4.78a | 4.61b | 4.55c | 3.90d | ** | ** | ** |
| Yeast | 4.78a | 4.08d | 4.35c | 4.52b | 4.97a | 4.25c | 4.70b | 4.63b | ** | ** | * |
| Mold | ND | ND | ND | ND | ND | ND | ND | ND | ND | ND | ND |
| Fermentation quality |  |  |  |  |  |  |  |  |  |  |  |
| pH | 4.67a | 4.37c | 4.58b | 4.56b | 4.90b | 4.61d | 4.72c | 5.06a | ** | ** | * |
| Lactic acid (% of DM) | 2.95b | 3.44a | 3.30a | 3.34a | 1.85a | 1.70b | 1.71b | 1.62c | ** | ** | NS |
| Acetic acid (% of DM) | 0.36a | 0.31b | 0.35a | 0.30b | 0.64b | 0.59b | 0.60b | 0.71a | NS | ** | NS |
| Propionic acid (% of DM) | ND | ND | ND | ND | ND | ND | ND | ND | ND | ND | ND |
| Butyric acid (% of DM) | 0.53a | ND | 0.44b | 0.49b | 0.96a | 0.86b | 0.80b | 0.97a | ** | ** | ** |
| NH<sub>3</sub>-N (g/kg of DM) | 0.37a | 0.21b | 0.28b | 0.20b | 0.66a | 0.59b | 0.52c | 0.61b | ** | ** | ** |
| Chemical composition |  |  |  |  |  |  |  |  |  |  |  |
| DM (%) | 17.48 | 17.82 | 16.75 | 16.53 | 24.00 | 23.99 | 22.90 | 23.02 | ** | NS | NS |
| OM (% of DM) | 90.21 | 90.23 | 90.37 | 90.51 | 90.10 | 90.32 | 90.06 | 90.31 | NS | NS | NS |
| CP (% of DM) | 29.22 | 29.48 | 30.04 | 29.31 | 26.72 | 27.06 | 26.62 | 26.39 | ** | NS | NS |
| NDF (% of DM) | 37.34a | 37.18ab | 36.24c | 36.53bc | 38.20a | 38.91a | 38.44b | 38.36b | * | ** | NS |
| ADF (% of DM) | 26.44a | 26.21a | 23.85b | 23.06c | 26.57b | 26.40b | 26.74a | 26.47b | ** | ** | ** |
| ADL (% of DM) | 6.90c | 7.05b | 7.37a | 6.93bc | 7.10b | 7.04b | 7.49a | 7.52a | ** | ** | ** |
| WSC (% of DM) | 6.32a | 4.00c | 6.30a | 5.09b | 6.21a | 3.83c | 5.18b | 5.02b | ** | ** | NS |
| IVDMD (%) | 58.41b | 60.54a | 59.27b | 55.67b | 58.47b | 60.11a | 57.72ab | 55.60b | * | * | NS |
| Gas production (ml) | 62.00b | 63.47a | 60.07b | 61.07b | 61.40a | 59.57a | 60.67ab | 58.20b | * | * | NS |

<sup>a-d</sup> Data are means of three samples, means of inoculation treatments within a row with different superscripts differ. \* *P* < 0.05; \*\* *P* < 0.01.
PM, paper mulberry; WPM, wilted paper mulberry; CH, *Lactobacillus plantarum*; AC, *Acremonium cellulase*; CH + AC, combination of CH and AC; S, silage; A, additive; S × A, interaction between silage and additive; cfu, colony-forming unit; FM, fresh matter; ND, not detected; DM, dry matter; NH<sub>3</sub>-N, ammonia nitrogen; OM, organic matter; CP, crude protein; NDF, neutral detergent fiber; ADF, acid detergent fiber; ADL, acid detergent lignin; WSC, water-soluble carbohydrates; IVDMD, in vitro dry matter digestibility; NS, not significant.

The PM silages had good fermentation pattern, with higher (*P* < 0.05) lactic acid contents and lower (*P* < 0.05) pH, butyric acid, and ammonia nitrogen (NH_3_-N) contents than those of the WPM silages. The CH, AC, and CH + AC-inoculated PM silages had higher (*P* < 0.05) lactic acid content and lower (*P* < 0.05) butyric acid content than those of control silage. The S and A influenced (*P* < 0.05) silage pH, lactic acid, and NH_3_-N contents, while the acetic acid were not. The A also influenced (*P* < 0.05) the acetic acid content. S × A influenced pH and NH_3_-N (*P* < 0.05), while the other fermentation product were not. After 60 days of ensiling, the OM contents of the PM and WPM silages were ranging from 90.21 to 90.51% of DM. Compared to WPM silages, the DM, NDF, ADF, and ADL contents of PM silages were lower (*P* < 0.05), while the contents of CP and WSC were higher (*P* < 0.05). The NDF and ADF contents of the AC and CH + AC-treated silages were lower (*P* < 0.05) than those of control and CH-treated silages in PM and WPM. The contents of DM, OM, and CP in the all PM and WPM silages did not show marked differences. S influenced (*P* < 0.05) DM, CP, NDF, ADF, ADL, WSC, and contents, but the OM did not. A influenced (*P* < 0.05) NDF, ADF, ADL, and WSC contents, while not the DM, OM, and CP. The interaction between S and A (S × A) influenced (*P* < 0.05) ADF and ADL contents, but the other chemical composition did not. In the PM, the IVDMD and gas production of CH-treated silage at 72 h incubation were higher (*P* < 0.05) than other silages. In the WPM, the IVDMD and gas production of control silage at 72 h incubation were higher (*P* < 0.05) than other silages. S and A influenced (*P* < 0.05) IVDMD and gas production, but the interaction between S and A (S × A) influenced (*P* < 0.05) did not.

### Correlation analysis of the bacterial community with fermentation products

Correlation analysis between the bacterial community and terminal fermentation products and correction networks using the Python tool at genus and species levels are shown in Fig. 5. In Fig. 5A, the moisture, LAB, pH, lactate, acetate, and NH_3_-N all had highly positive correlated (*P* < 0.01) with genus *Stenotrophomonas*. In addition, the content of lactic acid and LAB were highly positive correlated (*P* < 0.05) with *Lactobacillus* and *Pediococcus*. In Fig. 5B, the average maximum abundance was from the genus *Lactobacillus*, which positively correlated with *Exiguobacterium* (0.51). *Enterobacter* was the second-most abundant genus, which was negatively correlated with the genus *Pantoea* (0.49) and *Weissella* (0.42).

**FIG 5.**
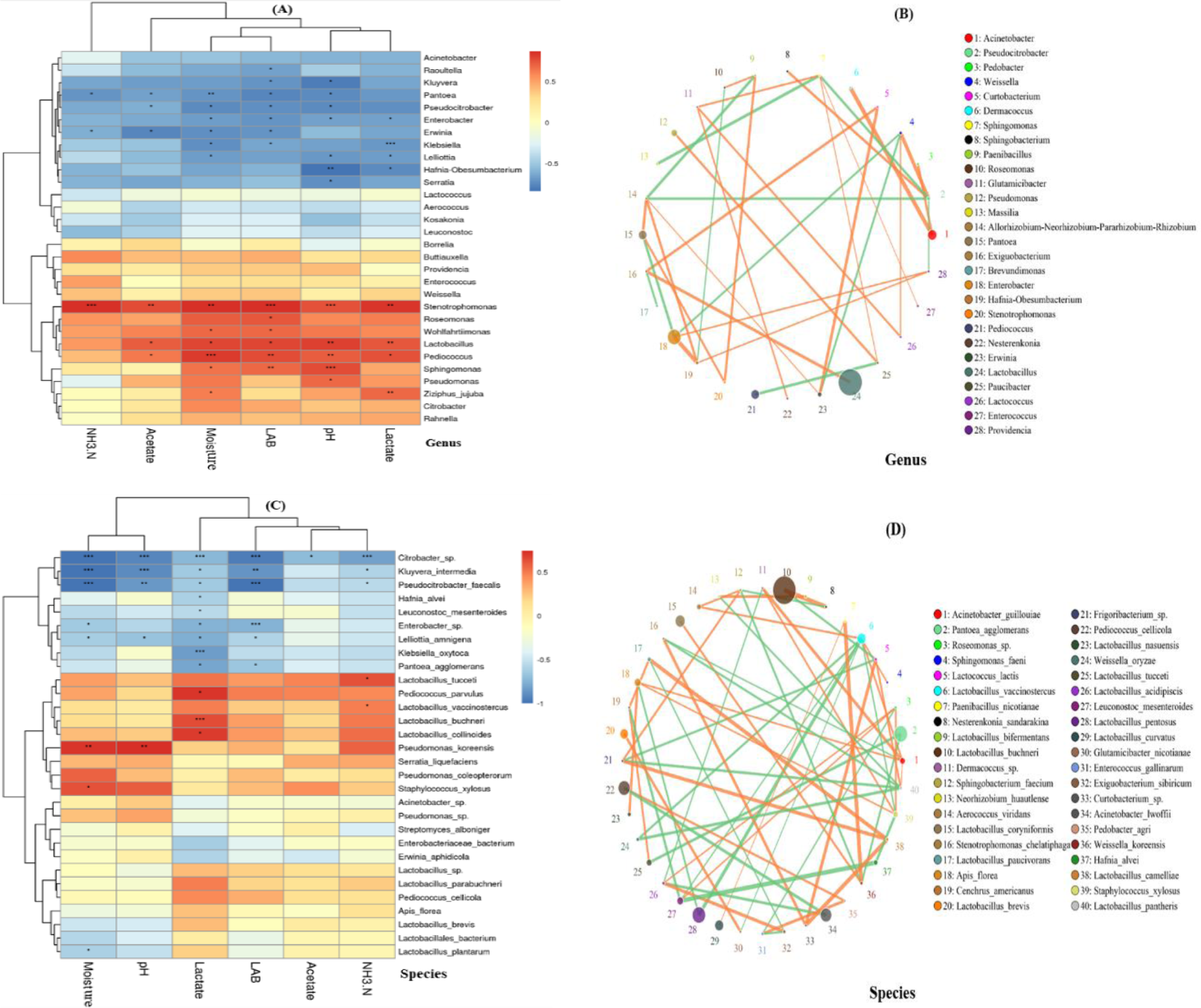
Correlation analysis (A and C) between bacterial community and terminal fermentation products and correction networks (B and D) using Python tool at genus and species levels. The Fig. 5A and 5B showed the correction analysis of the bacterial community with moisture, LAB, pH, lactate, acetate, and NH_3_-N at genus and species levels, respectively. Moisture, LAB, organic acid, and NH_3_-N information are displayed horizontally, respectively, and the bacterial community information is displayed vertically. The corresponding value of the middle heat map is the Spearman correlation coefficient r, which ranges between −1 and 1, r < 0 indicates a negative correlation (blue), r > 0 indicates a positive correlation (red), and ‘*’, ‘**’and ‘***’ represent *P* < 0.05, *P* < 0.01, and *P* < 0.001, respectively. The Fig. 5C and 5D showed the correction networks among microorganisms at genus and species levels, respectively. The circle represents the species, the circle size represents the average abundance of the species, the line represents the correlation between the two species, the thickness of the line represents the strength of the correlation, and the colour of the line: orange represents positive correlation, green means negative correlation.

In Fig. 5C, the moisture had highly positive correlated (*P* < 0.05) with species *Pseudomonas koreensis* (0.72) and *Staphylococcus xylosus*. The pH had highly positive correlated (*P* < 0.01) with species *Pseudomonas koreensis*. The NH_3_-N had the positive correlated (*P* < 0.01) with *Lactobacillus tucceti* and *Lactobacillus vaccinostercus*. In Fig. 5D, the most abundant were *Lactobacillus buchneri*, which was highly positive correlated with *Nesterenkonia sandarakina*. The second abundant was *Lactobacillus pentosus*, which was highly negative correlated with *Lactobacillus vaccinostercus*. The third abundant was *Pantoea agglomerans*, which was highly negative correlated with *Lactobacillus tucceti*.

### The impacted metabolic KEGG Pathways in PM and WPM before and after ensiling

After assigning orthologous sequences in the metagenome based on Kyoto Encyclopedia of Genes and Genomes (KEGG) gene database and pathway database, a total of 46 metabolic categories were obtained. All the KEGG metabolic pathways showed that, the global and overview maps, carbohydrate metabolism pathway, and amino acid metabolism pathways became the predominant metabolic categories in the raw materials and their silages of PM (Fig. 6A) and WPM (Fig. 6B). The global and overview maps of the most predominant metabolic categories included metabolic pathways (sequence ID 01100) for biosynthesis of secondary metabolites (01110), microbial metabolism in diverse environments (01120), carbon metabolism (01200), 2-oxocarboxylic acid metabolism (01210), fatty acid metabolism (01212), biosynthesis of amino acids (01230), and degradation of aromatic compounds (01220). PM had a higher number (*P* < 0.001) of metabolic pathways in silage than in the raw material, while WPM displayed the opposite. Higher (*P* < 0.001) carbohydrate metabolism and lower (*P* < 0.001) amino acid metabolism pathways were found in the PM and WPM silages than in their raw materials.

**FIG 6.**
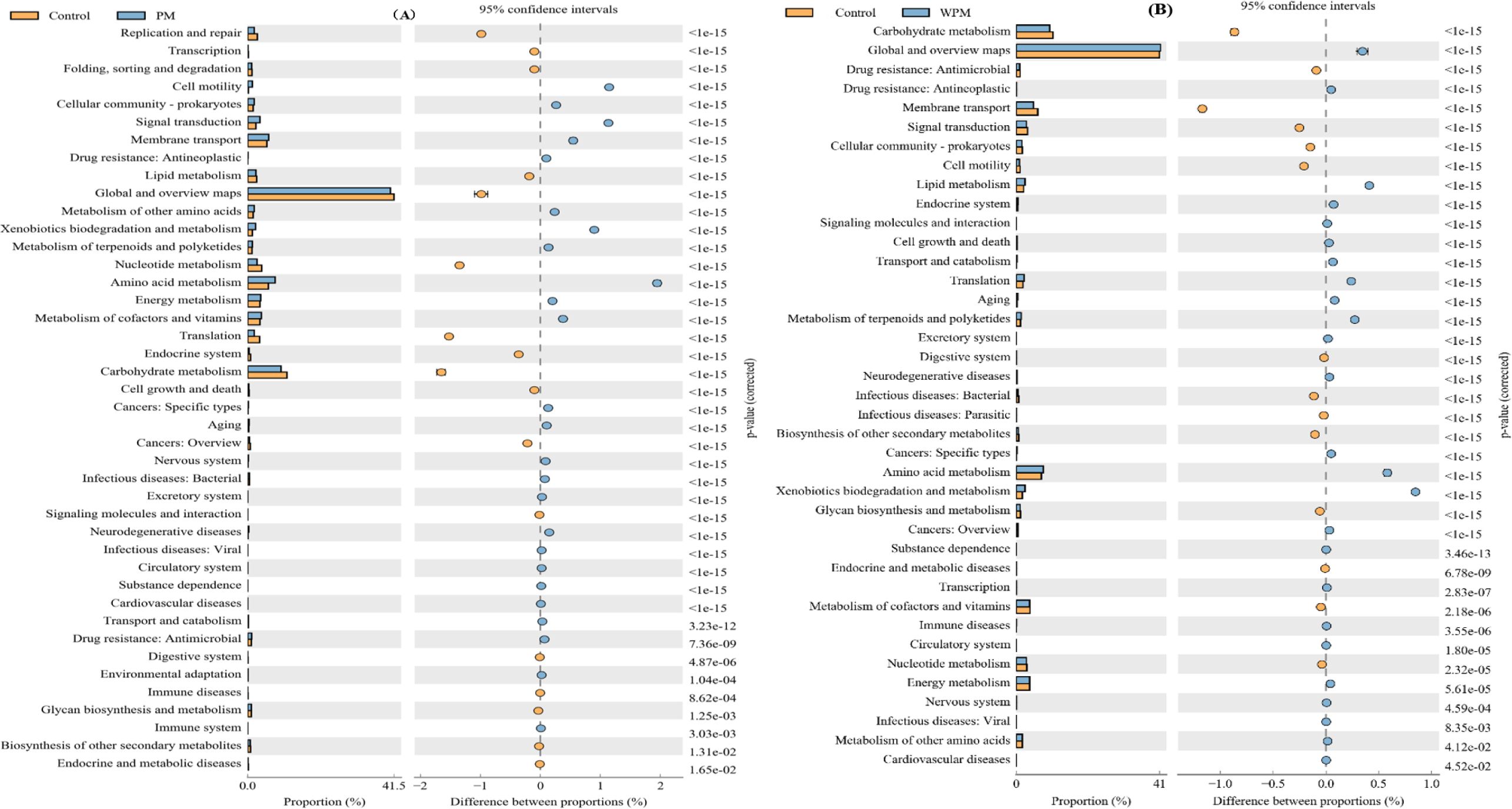
The impacted KEGG metabolic pathways on the second level in PM and WPM and their silages. KEGG, Kyoto Encyclopedia of Genes and Genomes; PM, paper mulberry; WPM, wilted paper mulberry

## DISCUSSION

Alpha diversity reflects the bacterial abundance and species diversity of a single sample. Chao1 and ACE indexes measure bacterial abundance, and Shannon and Simpson indexes are used to measure species diversity. In the present study (Table 1), the alpha diversity of PM were lower than those in WPM before ensiling. The local humid and hot environment during wilting process may lead to the increase of harmful microorganisms in the WPM. After ensiling, the Shannon and Simpson indexes were lower than those in raw materials of PM and WPM. Generally, after silage fermentation, the complex microbial communities of the raw materials are gradually replaced by LAB, and microbial diversity will sharply reduce, which is one of the criteria for the successful silage (8). In this study, after ensiling, compared to the control silage, the CH-inoculated PM silages had lower Shannon and Simpson indexes, while the WPM silages were opposite. The possible reason is that the LAB addition inhibited the growth of other microorganisms, result in reduce species diversity. On the other hand, during the wilting process, the excessive reproduction of microorganisms consumed the WSC as fermentation substrate, thereby weakened the role of LAB during fermentation, and increased the species diversity in WPM silage.

The taxa shared by all individuals in each group were deemed to represent core bacterial communities. As shown in Fig. 1, the diversity of the bacterial core microbiome (composed of 80 shared OTUs in silages) declined after ensiling in the two control silages compared to their respective raw materials. Before ensiling, the epiphytic microorganisms on the surface of the woody plants displayed high diversity. After silage fermentation, the LAB produced lactic acid which reduced the pH, thus inhibiting the growth of microbes and reducing the bacterial diversity. LAB addition yielded the lowest unique OTUs in the PM silage but did not greatly reduce the OTUs in the WPM silage. For the PM silages, the lower diversity was likely because the relatively lower pH values in LAB-inoculated silages can limit the growth of harmful microbes in the acidic environments (9). In addition, the exogenous LAB can rapidly proliferate to become dominant species in the bacterial communities, which in combination with silage acidification and antagonistic activity towards other bacteria, led to a decline in bacterial diversity, as well as improved silage quality (10). The absence of a reduction in OTUs in the WPM silages could be due to the inability of the LAB used in this experiment to adapt well to the wilted environment where the proliferation of undesirable microorganisms and the nutrient consumption makes the LAB unable to dominate the silage fermentation. These results indicate that the similar microbial communities in successful silage not only share core bacteria but may also display a reduction in unique bacteria.

The diversity, function, and structure of microbial community are hot microbial ecology topics and have been conducted researches in many media (11). As far as we know, this is the first report on the bacterial community of PM silage. It was demonstrated that the *Pseudomonas* is one of the most common bacterium in soil, and it can survive in an anaerobic environment (12), and the negative correlation between *Pseudomonas* and NH_3_-N concentration and yeast count in silages, indicating that the presence of *Pseudomonas* might contribute to protein preservation and yeast repression (13). In this study, as shown in Fig. 2, *Pseudomonas* was the most abundant genus in PM raw material, this bacteria may retain the nutrients of PM as direct-cut fermented silages, the higher CP and lower NH_3_-N, and yeast counts (Table 3) supported this theory.

In the wilting process, *Pseudomonas* spp. cannot survive, and harmful microorganisms multiply. The increased *Acinetobacter* abundance may have resulted to the acetic acid production in WPM silage. The LAB genus *Lactobacillus, Lactococcus, Pediococcus*, and *Enterococcus* are desirable functional bacteria during the ensiling, and they have been widely utilized to improve the fermentation quality of silage (14). Generally, lactic acid producing cocci is the dominant LAB and initiate lactic acid fermentation during the early stages of ensiling, as fermentation process and the pH decreases, lactobacilli will grow rapidly to reach a high abundance (11). This phenomenon suggested that the *Lactobacillus* and *Enterococcus* were the important genus for silage fermentation, and that the LAB inoculant can change the community structure and genus abundance during ensiling.

As shown in Fig. 3, in the PM and WPM raw material, the dominant species were *Pantoea agglomerans* and *Acinetobacter* spp., and the *L. plantarum* numbers were very low. The results demonstrated that similar to forage, the woody plant usually has a low abundance of *Lactobacillus* species and a high abundance of undesirable bacteria before ensiling. Silage fermentation is an anaerobic, closed, solid-fermentation system, and a higher relative abundance of *Lactobacillus* species was frequently observed in this system, and the different species and proportions of lactobacilli had different effects on the quality of silages (15). In the present study, the most abundant microorganisms in PM silages after fermentation were *Lactobacillus plantarum*, especially in the CH-treated silage (96.92%). *Lactobacillus plantarum* can act as a homofermentative LAB and grow well in an acidic environment. Silage inoculation with this strain will promote lactic acid fermentation. The relatively abundant microbe species in the WPM silages were *Enterobacter* spp. (17.43–46.64%), which are one of the principal competitors of LAB for available WSC in silages and produce acetic acid. This could explain why the lactic acid levels (1.62–1.85% of DM) in PM silage are usually lower than the standard concentration in high-quality silage, which was determined to be caused by poor fermentation. The lower abundances of all *Lactobacillus* species and the higher abundances of undesirable bacterial species in the untreated PM and WPM silages may explain their poor quality.

In PM and WPM raw materials (Fig. 4), there were three phyla including *Proteobacteria, Firmicutes*, and *Actinobacteria. Proteobacteria* is a major phylum belonging to Gram-negative bacteria, that include a wide variety of pathogenic genera, such as *Escherichia, Salmonella, Vibrio, Helicobacter, Yersinia, Legionellales*, and many others. In this study, *Proteobacteria* was the most abundant phylum in the PM and WPM, exceeding 80% in raw materials. After ensiling, *Proteobacteria* decreased while Gram-positive *Firmicutes* increased rapidly and became the dominant phylum. The silage fermented related bacteria such as Bacillus, Clostridium and all genus of LAB were belonged to this phylum. Therefore, the bacterial community during silage fermentation displayed a dynamic change from Gram-negative bacteria to Gram-positive bacteria, indicating that the abundant pathogenic bacteria was presented in raw materials, after ensiling, the LAB that are beneficial to silage fermentation were converted into the dominant bacteria.

The fresh stem and leaves of PM have high nutrition value, and adequate balance of fiber, lipids, amino acid, vitamins, and minerals. For ensiling, LBC, WSC, moisture, and LAB of raw material are important factors affect silage fermentation. In the present study (Table 2), the WSC content of PM and WPM raw materials were 8.26 and 8.40% of DM, respectively, which are higher than the theoretical requirement (6–8% DM) (8), forecasting this woody plant can prepare as well-preserved silage. In this study, the moisture of PM and WPM raw materials were high (approximately 80%), after wilting, the moisture (73%) was close to the ideal level for silage preparation (16). The low counts of LAB (10^4^ cfu/g of FM) and higher amounts of undesirable microorganisms (10^3^–10^5^ cfu/g of FM) were observed in the PM and WPM raw material, which may lead to poor quality in natural fermentation. The CP and effective protein contents of PM were 29% of DM and 93% of CP, which were much higher than alfalfa (17). The higher proportion of effective protein means better nutritional value of the protein given, and the non-protein is less efficient in utilization for ruminants. In addition, the NDF and ADF contents of PM raw material were low than alfalfa and other forages. The lower NDF and ADF contents resulted in higher IVDMD and digestible energy (DE). The gas production content in PM higher than WPM, due to the well preserved silage. The calcium, phosphorous, magnesium, and potassium contents in the PM raw material were higher than other forages. Therefore, the PM can be used as a superior high protein and digestible woody forage resources for ruminants.

As shown in Table 3, the LAB and yeast counts increased, but the aerobic bacterial counts decreased, and the mould and coliform bacteria were below the detection level (10^2^ cfu/g of FM) in all the silages. These results showed that the woody PM plant can be prepared as good quality silage similar to other forage crops. In this study, the PM silages had better fermentation quality than the WPM silages, especially in the CH-silage, with lower pH and NH_3_-N content, and higher lactic acid content. The inoculant strain, *L. plantarum*, belongs to the homofermentative LAB, which can inhibit the growth of *Clostridia* and decrease the content of NH_3_-N, which is an indicator of high-quality fermentation (2). In the WPM, the rapid growth of harmful microbes during the wilting process results in poor silage fermentation. The microbial community and abundance analysis in this study also confirmed this point.

The AC and CH+AC-treated PM silages had markedly lower NDF and ADF contents compared with other silages. The cellulases, mainly endoglucanases, cellobiohydrolases, and β-glucosidases, catalyse the hydrolysis of cellulose and any mixture or complex of such enzymes that acts serially or synergistically can be utilised to decompose cellulosic material. The cellulases produced by *A. cellulolyticus* in this study contain glucanase and pectinase, indicating that this cellulase should be able to degrade cellulose effectively. However, the addition of cellulase did not affect the NDF and ADF of the WPM silage, and the WSC content decreased. The reason may be that the rapid multiplication of harmful microbes depleted sugars in the wilting process, which prevented the growth of LAB and the enzyme activity during ensiling.

Microorganisms affect the silage quality via a series of metabolites, e.g. *Lactobacillus* mainly affects lactic acid production, while *Enterobacteria* can ferment lactic acid to acetic acid and other products (18). Correlation analysis between the bacterial community and terminal fermentation products and correction networks using the Python tool at genus levels are shown in Fig. 5. In this study, we also performed a correlation analysis of the composition of the microorganism population with the contents of fermentation products after ensiling. The lactic acid content and LAB count were positively correlated with the genera *Lactobacillus* (0.78 and 0.73, respectively) and *Pediococcus* (0.78 and 0.76, respectively). These results were consistent with previous reports (18), indicating that the LAB genus plays an important role in silage fermentation. In Fig. 5B, the average maximum abundance was from the genus *Lactobacillus*, which positively correlated with *Exiguobacterium* (0.51). *Exiguobacterium spp.* are globally diverse organisms that are found in a variety of environments, and some species, in addition to dynamic thermal adaption, are also halotolerant and can grow within a wide range of pH values. This characteristic confirmed that *Exiguobacterium* may grow in a silage environment, perhaps impacting the silage fermentation together with *Lactobacillus. Enterobacter* was the second-most abundant genus, which was negatively correlated with the genera *Pantoea* (0.49) and *Weissella* (0.42), showing that there is a distinct relationship between acetic acid bacteria and the production of acetic acid in silage.

As shown in Fig. 5C, the moisture content had a high positive correlation with population counts of *P. koreensis* and *Staphylococcus xylosus*. The results suggest that the moisture content affected the community of Gram-negative *Pseudomonas* and Gram-positive *Staphylococcus* species, with lactic acid fermentation and pH reduction potentially inhibiting the growth of these bacteria. Generally, LAB fermentation can effectively inhibit protein decomposition, but some LAB in this experiment had the opposite effect. *Lactobacillus plantarum* showed a positive correlation with the lactic acid concentration (0.51) and a negative correlation with the pH (0.53), suggesting this bacterium can grow under low pH conditions, and play an important role in silage fermentation. In this study, *Lactobacillus buchneri* was associated with higher DM loss. This can be explained in part by its being a heterofermentative LAB. In heterofermentative fermentation, an equivalent molecule of carbon dioxide is generated for every molecule of acetic acid formed; therefore, a considerable loss of DM would be expected with heterofermentative fermentation in silage (8). As shown in Fig. 5D, the most abundant species was *L. buchneri*, which positively correlated with *Nesterenkonia sandarakina* (0.70). *Lactobacillus buchneri* is a heterofermentative LAB that produces acetic acid to increase the aerobic stability of silage. *Nesterenkonia sandarakina* is a halophilic actinobacterium that produces an antibacterial substance (5). During silage fermentation, this bacterium may inhibit the growth of harmful microorganisms together with *Lactobacillus spp*. The second most abundant species was *Lactobacillus pentosus* and its population negatively correlated with *L. vaccinostercus* (0.58). *Lactobacillus pentosus* is recognised to play an important role in olive fermentation. *Lactobacillus pentosus* is closely related to *L. plantarum* based on its physiological and biochemical characteristics, therefore this LAB may be able to limit the growth of other LAB such as *L. vaccinostercus.* The third abundant species was *P. agglomerans*, which negatively correlated with *L. tucceti* (0.46). *Pantoea agglomerans* is a ubiquitous bacterium commonly isolated from plant surfaces, seeds, fruit, and animal or human faeces. *Lactobacillus tucceti* is usually isolated from sausage. Generally, these bacteria cannot be isolated from silage fermentation in forage crops and grasses, and this is the first report of their isolation from woody plant silage prepared from PM. We are interested in studying their true function in silage fermentation in future studies.

Metabolic gene clusters or biosynthetic gene clusters are tightly linked sets of mostly non-homologous genes participating in a common, discrete metabolic pathway. The genes are in physical proximity to each other in the genome, and their expression is often coregulated (19). Metabolic gene clusters are also involved in nutrient acquisition, toxin degradation, antimicrobial resistance, and vitamin biosynthesis (20). These metabolic genes affect the microbial communities and silage fermentation. PM has high nutritional value and contains a variety of biologically active ingredients, which makes it a potential animal feed resource with economic and practical value. More than 40 flavonoids and terpenes with antioxidant and anti-inflammatory properties (21) were found in PM. The contents of flavonoids, polyphenols, fructose, and other constituents in PM are higher than in many other plants. The flavonoids are a major class of secondary plant metabolites in the plant kingdom with many functions such as pigmentation and antimicrobial activity (22), which plays an important role in disease prevention (23). In general, most of the metabolic categories increased over time, indicating that the metabolic intensity of microbial communities increased with the advance of microbial diversity and abundance (24). All the KEGG metabolic pathways, global and overview maps, carbohydrate metabolism pathways, and amino acid metabolism pathways became the predominant metabolic categories in the PM and WPM raw materials and their silages. This indicated that three metabolic categories were of key importance for microbial metabolism. In this study, the high incidence of amino acid metabolic pathways displayed an ability to degrade the macromolecular proteins in PM into amino acids or peptide substances easily absorbed by the body, with some metabolites possibly having antibacterial activity, greatly enhancing the functionality of PM. The metabolic pathways of pyruvate metabolism, glycolysis, and butanoate metabolism occupied a dominant position in carbohydrate metabolism, which might have a significant influence upon the taste (sour and sweet) and palatability of PM silages for animals. We found that the PM and WPM silages had greater (*P* < 0.001) digestive capacity than their raw materials, indicating the silage may improve feed digestibility and livestock production capacity. Thus they have the potential to partially replace expensive supplements in cattle feed, as they can promote animal growth without any damaging effects on animal health. Therefore, the development of this unconventional woody feed is of great significance for animal production.

In conclusion, the community structure, species diversity, and metabolic gene cluster of microbes related PM silage fermentation were studied by PacBio SMRT. The microbial diversity was higher in raw materials than that of silages, and the microbial additives reduced the bacterial diversity and improved PM silage quality. Silage fermentation process displayed the dynamic shift of dominant bacteria from Gram-negative to positive, lead to the LAB became the most dominant genus and species in silages. The PacBio SMRT reveals more specific microbial information of affect to silage fermentation, and the carbohydrate and amino acid metabolism were the important metabolic pathways in PM silages.

## MATERIALS AND METHODS

### Woody plants and silage fermentation

The experiment was conducted on 22 May 2019 at a mountain experimental station (106°37′E, longitude, and 25°73′N, latitude) of Guizhou University in Changshun, Guizhou, China. The climate of the station is characterised as a mid-subtropical monsoon humid climate zone, with an average annual temperature of 15.5°C, relative humidity of 81%, and annual rainfall of 1250–1400 mm. There is no severe cold during winter and no extreme heat during summer, and the annual minimum temperature is -15.5°C and the extreme maximum temperature is 40.7°C. The PM (Hybrid *B. papyrifera*-Zhongke No. 1) plants cultivated at the experimental station were approximately 1.5 m high and were harvested 4–5 times a year. The fresh stems and leaves were obtained at the juvenile stage from the first cutting with three field replications leaving stubble of 15 cm.

After harvest, the fresh stems and leaves were immediately cut into approximately 1–2 cm lengths using a chopper (92–2S, Sida Agri-Machine Co., Ltd, Luoyang, China). The chopped materials were divided into three fractions. The first fraction was collected as the fresh samples into sterilised bags that were kept in iceboxes, then immediately transported to a laboratory to determine the chemical composition. The second fraction was destined for silage making. To study the effect of moisture adjustment on silage fermentation, 50% of the fresh samples were wilted for 6 h in the shade on a black plastic sheet on a concrete floor. The WPM was turned twice at 3-h intervals. The third fraction of the material was used for fresh silage preparation.

Two kinds of microbial inoculants were used as silage additives according to the commercial manufacturer’s guidelines. The experimental treatments for PM and WPM were designed as 1) a control (without inoculant), 2) a treatment with the addition of the CH, 3) a treatment with the addition of AC, and 4) a treatment with the addition of a combination of LAB and cellulase enzyme (CH + AC). The CH inoculum size was 5 mg/kg of power as 1.0 × 10^5^ cfu/g of FM. AC is produced by *Acremonium cellulolyticus*, and contains glucanase, pectinase, and carboxymethyl-cellulase activity at a total cellulase activity of 7,350 U/g. AC cellulase was added at 0.01% of FM. Both of the treatments with additives (CH and AC) were sprayed with deionised water using an electronic sprayer (SSP-5H, Fujiwara Sangyo Co., Ltd, Miki, Japan). The control treatment was sprayed with the same amount of deionised water.

Three replicates of silage were prepared with a laboratory fermenter. After thoroughly mixing the materials with additives, approximately 35 kg of PM material (fresh and wilted) for each replicate was packed into 30 L polyethylene drum silos (260 mm, Huafang Plastic Co., Ltd, Shandong, China). All the silos were then compacted to exclude air and stored at ambient temperature (25°C to 38°C). After 60 days of ensiling, the silos were opened, and triplicate samples of each treatment were divided into three portions: the fresh portion was used for microbial community analysis; the dried portion was used for the analysis of its chemical composition, and the extracted liquid portion was used for the analysis of fermentation characteristics.

### Analysis of chemical composition, energy, and macro minerals

The pre-ensiled materials and silage samples of PM and WPM were dried in an oven for 48 h at 65°C until they reached a constant mass. After drying, the samples were ground in a high-speed vibrating sample mill (model T1-200, for use with two containers of 50 ml working capacity each, CMT Co., Ltd, Japan). The chemical and protein composition, energy content, and macro mineral composition were analysed as described by Cai (2). The LBC was determined as described by Muck (25). The WSC was determined using a high-performance liquid chromatography (HPLC) system (LC-2000 plus, JASCO Corporation, Tokyo, Japan) as described by Cai (26).

### Analysis of silage fermentation

The final fermentation products of the PM and WPM silages were analysed using the cold-water extract method as described by Cai (26). The remaining wet silage sample (10 g) was homogenised in 90 mL of sterilised distilled water and kept in a refrigerator at 4°C for 24 h. Thereafter, the extract samples were filtered through quantitative ashless filter paper (circle size: 5A, 110 mm; Advantec Co., Ltd, Tokyo, Japan). The pH was measured using a glass electrode pH meter (D-71, Horiba Co., Ltd, Kyoto, Japan). The ammonia nitrogen content of the silages was determined by the steam distillation of the filtrates as described by Cai (26) and using the Kjeltech auto distillation system (2200, Foss Tecator, Hoganas, Sweden). The silage filtrates were shaken with cation exchange resin (Amberlite, IR 120B H AG; Organo Corporation, Tokyo, Japan) and centrifuged at 6,500 × *g*, at 4°C for 5 min. The supernatants were passed through a 0.45 µm filter under pressure, and the filtrates were then injected into an HPLC system (LC-2000 plus, JASCO Corporation, Tokyo, Japan) to determine the organic acid contents following the methods described by Cai (26).

### In vitro gas production and animal care

The in vitro gas production in ruminants was determined using calibrated glass syringes in a randomised experimental design. The rumen fluid was collected from three healthy mature Hainan black goats with a stomach tube sucker before morning feeding for measurement of IVDMD and gas production. The rumen fluid was strained through gauze and mixed with a buffer solution as described by Menke and Steingass (27). The use of an animal procedure in this study was approved by the Institutional Animal Care and Use Committee (IACUC) at China Agricultural University (DK 996 No. 20144-2), Beijing, China.

### Microbial population analysis

The microbial population of silages was measured using a plate count method as described by Cai (10). Samples (10 g) were blended with 90 mL of sterilised saline solution (8.50 g/L NaCl) and homogenised for 5 min in a Stomacher lab blender (400, Seward, UK). The resulting suspension was serially diluted from 10^−1^ to 10^−8^ using a saline solution. A 0.05 mL aliquot from each diluted suspension was spread on agar plates. LAB were cultured on Lactobacilli de Man, Rogosa, and Sharpe agar medium (Difco Laboratories, Detroit, MI, USA) in an anaerobic box (TEHER Hard Anaerobox, ANX-1; Hirosawa Ltd, Tokyo, Japan) and colonies were counted. Aerobic bacteria were cultured on nutrient agar medium (Nissui-Seiyaku Co., Ltd, Tokyo, Japan) under aerobic conditions. Coliform bacteria, which can be distinguished from other bacteria by the blue colour of their colonies, were cultured and counted using blue light broth agar (Nissui-Seiyaku Co., Ltd, Tokyo, Japan). Yeasts and moulds were counted on potato dextrose agar medium (Nissui-Seiyaku Co., Ltd, Tokyo, Japan) with a sterilised tartaric acid solution (pH 3.5). All plates were incubated at 30°C for 2–3 days and the microbial colonies were reported as viable numbers of microorganisms in cfu/g of FM.

### Bacterial community analysis: DNA extraction and SMRT sequencing

Each sample (25 g) was frozen (−80°C) and passed through a 4 mm sieve after freeze-drying and smashing, and a subsample (5 g) was ball-milled for 1 min at room temperature. Total genomic DNA from each sample was extracted using the TIANamp Bacteria DNA isolation kit (DP302-02, Tiangen, Beijing, China). The quality of the extracted DNA was monitored on 1% agarose gels using electrophoresis and spectrophotometry (optical density at 260/280 nm ratio). All the DNA samples were purified through a DNA kit column (DP214-02, Tiangen, Beijing, China) and kept at −20°C until further analysis (6). The SMRT sequencing was performed on a PacBio RS II instrument (Pacific Biosciences, Menlo Park, CA, USA) using P6-C4 chemistry (7).

### Sequence analyses

Raw data were processed using the protocol RS_Readsofinsert.1 in SMRT Portal version 2.7 software (PacBio). Low-quality sequences were removed using the Quantitative Insights Into Microbial Ecology (QIIME) package (version 1.7). Then the extracted high-quality sequences, under 100% clustering of sequence identity, were aligned to obtain representative sequences using Python Nearest Alignment Space Termination (PyNAST) and Clustering and Classification Inference with U-Statistics (UCLUST) analyses (7). The unique sequence set was classified into OTU based on a 98.6% threshold identity using the UCLUST algorithm (7). Potential chimeric sequences in the representative set of OTUs were removed via the Chimera Slayer utility (28), and then the Ribosomal Database Project (RDP) II database classifier was used to classify different OTUs and annotate the taxonomic information for each OTU representative sequence based on Bergey’s taxonomy at the genus, family, order, class, and phylum levels, which classified at an 80% minimum bootstrap threshold (7). OTUs that occurred only once or twice were discarded.

The ACE, Chao1, Shannon, and Simpson indices were calculated using QIIME software to evaluate the alpha diversity (29). According to the analysis of the OTU clustering, and the research requirements to analyse both the common and unique information for different samples, we produced Venn diagrams using open source tools for R. Hierarchical cluster and heat map analyses were performed using the R-statistics tool. A phylogenetic circle tree based on sequences representative of the OTUs at the species level of all the samples was drafted using the Python tool. Correlation network analysis was also calculated using the Python tool. To analyse the functional profiles and differences in different groups, the metabolic potential of the microbial community and the composition of functional genes were postulated by assigning 16S rRNA marker gene sequences to functional annotations of sequenced metagenomic sequences based on the KEGG using the phylogenetic investigation of communities by reconstruction of unobserved states (PICRUSt), as described by Langille (30).

### Statistics analysis

Analysis of variance (ANOVA) was performed using the general linear model procedure of SAS software (rev. 9.4, Institute Inc., Cary, NC). The pH, LBC, microbial population, chemical composition, energy content, IVDMD, gas production, and macro mineral data for fresh forages were subjected to one-way ANOVA. The data for the microbial population, fermentation quality, chemical composition, IVDMD, and gas production of silages at 60 days were subjected to two-way ANOVA factorial arrangement, with forage and additive treatments (2 × 4) as the main variables. Statistical significance was declared at *P* < 0.05.

## ACKNOWLEDGEMENTS

This study was supported by National Key R&D projects ‘Processing technology research and demonstration of high-quality forage silage and molded product’ (2017YFD0502102), Ministry of Science and Technology, China, and JIRCAS Visiting Research Fellowship Program 2019-2020, Japan International Research Center for Agricultural Sciences, Japan.

